# A multidisciplinary approach for the study of age-dependent expression patterns in porcine experimental wounds

**DOI:** 10.1101/2025.11.04.686454

**Authors:** Cecilie Bækgård, Lars Andresen, Marie Høy Hansen, Henrik Elvang Jensen, Kristine Freude, Kristiane Barington

**Author notes:** Corresponding author. E-mail address (C. Bækgård).

## Abstract

Determining the age of wounds in both human and veterinary subjects is a critical aspect of forensic pathology. Gross and histopathological evaluations are used for age estimations, however, these evaluations are subjective as they rely on the opinion and experience of the examining pathologist. Therefore, the aim of the study was to create a multidisciplinary approach to discover accurate and objective indicators of wound age in veterinary forensic cases.

In the present study we utilized a porcine experimental wound healing model. Granulation tissue from wounds at different ages (5, 10, 15, 20, 25, 30, and 35 days) (n=188) and control skin samples (n=47) from 47 experimental pigs were evaluated macroscopically, histologically, immunohistochemically, combined with digital pathology, and flow cytometry.

Granulation tissue thickness peaked on day 10 post wounding. Full epithelialization was observed at the earliest on day 20 and the percentage of wound surface covered by epithelium increased with age. Leukocytes including hemosiderophages, neutrophils and accumulation of macrophages all displayed time-dependent infiltration into the wound bed. CD34 showed no time-dependent expression. CD105 expression peaked on day 15. CD45 showed a time-dependent expression pattern in the superficial part of the granulation tissue, with highest expression on day 5 and a decrease with increasing wound age.

The findings of this multidisciplinary approach indicate the potential to achieve accurate and objective wound age estimations in veterinary and human forensic investigations.

## Introduction

Obtaining an accurate estimation of wound age is of critical importance in both human and veterinary forensic investigations [1]. In cases of suspected violations of European and national animal protection legislation, the assessment of wound age may be used in a judicial process to determine the degree of neglect or assign responsibility [2, 3]. In Denmark, 79% of veterinary forensic post-mortem examinations involve pigs, and approximately one-third of these cases concern skin wounds [4, 5].

Currently, gross- and histopathological evaluations remain the approaches for determining wound age in veterinary forensic investigations [5, 6]. Granulation tissue thickness is used for age assessments, but in experimental wounds, thickness peaks on day 10 post wounding, after which it decreases [7]. Histopathological evaluation is used to assess the wound tissue for, e.g. the morphology of the granulation tissue, and inflammatory cell presence [7–9]. However, histopathological evaluation is somewhat subjective, as it relies on the opinion of the examining veterinary pathologist and their level of experience [10]. Often, wound ages are estimated simply as “several days”, “weeks”, or “months” [6]. To enhance objectivity, accuracy, and reproducibility of wound age assessments, we propose the combined use of immunohistochemistry (IHC) with digital cell detection and flow cytometry in addition to traditional methods. Flow cytometry allows for quantitative analysis of cell populations expressing markers of choice, while IHC enables spatial localization of specific markers within the wound tissue [11]. Digitalization of tissue slides followed by evaluation by tissue analysis software is hypothesized to provide a reproducible and objective method for estimating wound age.

The wound healing process is complex and highly regulated. It consists of four overlapping phases: hemostasis, inflammation, proliferation, and remodeling [12–16]. Mesenchymal and hematopoietic stem cells (MSCs and HCSs) contribute critically to wound healing through their regenerative and immunomodulatory functions. These bone-marrow derived cells reside in the skin during normal conditions and, upon injury, serve as a reservoir for a wide range of cell types that are essential during the healing process [17]. Multipotent MSCs are capable of differentiating into multiple cells lineages, such as adipocytes, fibroblasts, endothelial cells and keratinocytes [18, 19]. HSCs are capable of regenerating the entire population of hematopoietic cells (HCs), which include platelets, macrophages, neutrophils, and lymphocytes [20].

Stem cell populations are commonly identified based on expression of specific surface markers. MSCs are characterized by expression of cluster of differentiation 105 (CD105), whereas HSCs express markers such as cluster of differentiation 45 (CD45) and cluster of differentiation 34 (CD34) [17, 18]. CD105 is a marker of endothelial cells involved in angiogenesis [21–23]. CD45 is expressed on the surface of most HCs. A subset of these HCs, namely hematopoietic progenitor cells, also express CD34. These cells have been shown to aid in tissue repair by releasing factors that support angiogenesis, immune modulation, and tissue remodeling [24]. These cellular markers may change in the wound over time and could serve as potential biomarkers for estimating wound age.

A porcine experimental wound model offers opportunity to study mechanisms of the wound healing process. Additionally, the pig as an animal model offers significant anatomical, physiological and genetic similarities to humans. Porcine skin shares several commonalities to human skin, including comparable epidermal thickness, dermal collagen content, and immune response profiles [25–29].

This study characterizes time-dependent features of porcine granulation tissue by examining the morphological and cellular traits in experimentally induced wounds. Through a multidisciplinary approach, including gross and histological evaluation, IHC, digital cell quantification, and flow cytometry, the aim was to establish reliable indicators of wound age that may enhance the accuracy and objectivity of forensic age assessment of wounds.

## Materials and Methods

### Experimental wound model

In total, 50 female Yorkshire-Landrace crossbred pigs with bodyweights of 20-25 kg were acclimatized for one week. Each pig was housed individually with straw bedding and allocated into seven groups in the order they arrived at the facility. Two pigs died during anesthesia and one pig was euthanized within 12 hours after surgery due to respiratory distress. All three pigs were excluded from the study.

The experimental procedure was carried out as described by Barington et al [7] with minor alterations. Each pig was sedated using intramuscular injection of tiletamine and zolazepam (Zoletil 50 vet., Virbac, Carros, France) in combination with xylazine (Xysol vet., ScanVet, Fredensborg, Denmark), butorphanol (Butomidor vet., VetViva Richter GmbH, Wels, Austria), methadone (Comfortan vet., Dechra Veterinary Products, Uldum, Denmark), and ketamine (Ketador vet., Salfarm, Kolding, Denmark) before being prepared for sterile surgery. All pigs were washed with MediSkrub disinfecting soap, dried and disinfected with 85% alcohol. Pigs were put in sternal recumbency during surgery and received continuous intravenous propofol (Propofol “B. Braun”, 10mg/mL, B. Braun Medical, Melsungen, Germany) and fentanyl (Fentadon vet., Dechra Veterinary Products, Uldum, Denmark). On the back of the pigs, four surgical areas of 2 x 2 cm (locations A-D) were drawn located 4 cm lateral to the spine and 4 cm cranial and caudal to the last rib, respectively. Full thickness wounds (epidermis down to and including the subcutis) were created by incision (Figure 1). All wounds were left to heal by second intention for 5, 10, 15, 20, 25, 30, or 35 days. Sections of the skin excised from surgical area A were sampled as controls for each of the experimental approaches.

**Fig. 1.**
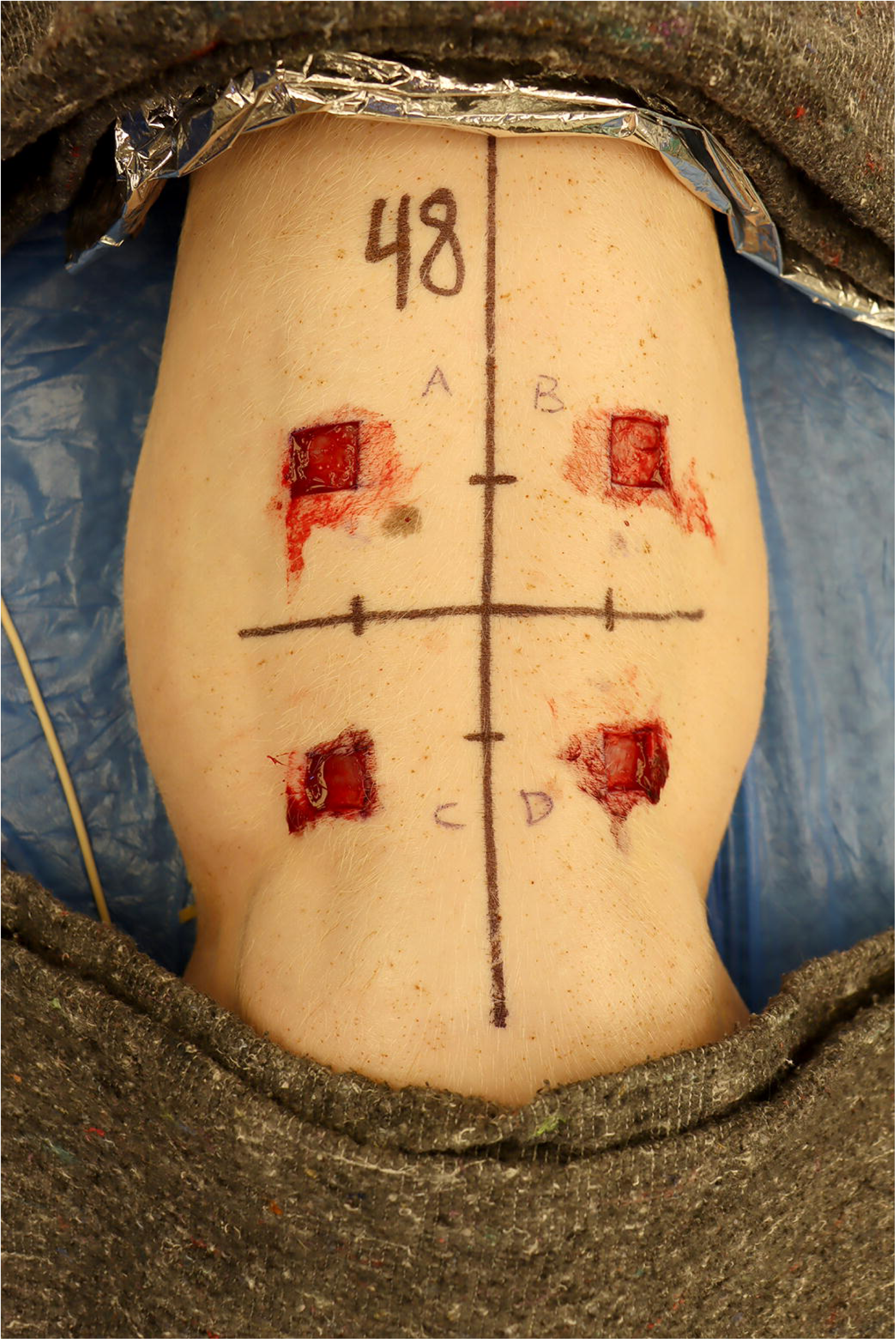
Anesthetized pig in ventral recumbency with full thickness wounds excised on the back. The wounds were located 4 cm lateral to the spine and 4 cm cranial (A and B) or caudal (C and D) to the last rib. A skin sample was taken as control from wound A

All pigs were given four injections of buprenorphine (Bupaq Multidose vet., 0.3 mg/mL, Salfarm, Kolding, Denmark) at 8-hour intervals. Upon indication of pain, additional pain treatment was administered.

At the timepoints mentioned above, the pigs were sedated using the same method as on the day of surgery, before being euthanized with intravenously administered pentobarbital (Exagon vet., Salfarm, Kolding, Denmark). Post-mortem, wounds A, B, C, and D were cross sectioned into two halves. From one half, a section with a thickness of approximately 5 mm was sampled for histological and IHC analysis. From the other half of wound A and D, a 0.5×0.5×0.5 cm sample of granulation tissue was taken from the center for flow cytometry.

### Gross evaluation

Granulation tissue was registered as present or absent. Additionally, the thickness of the granulation tissue was measured at both edges and at the center.

### Histology

Tissue samples were immersion-fixed in 10% neutral buffered formalin for 5-7 days, processed through graded concentrations of ethanol and xylene before being embedded in paraffin wax. From all wounds, tissue sections were cut at 4-5 µm and stained with hematoxylin-eosin (HE). All tissue sections were digitized using an Axioscan 7 with a 20x/0.8 objective (Zeiss, Germany). Scanned tissue sections were imported into QuPath v0.5.1 [30], and the granulation tissue was divided into a superficial and deep layer as previously described [5]. The granulation tissue was further divided into six approximately equal areas (wound areas I to VI) (Fig. 2). Histological assessment was carried out blinded to wound age.

**Fig. 2.**
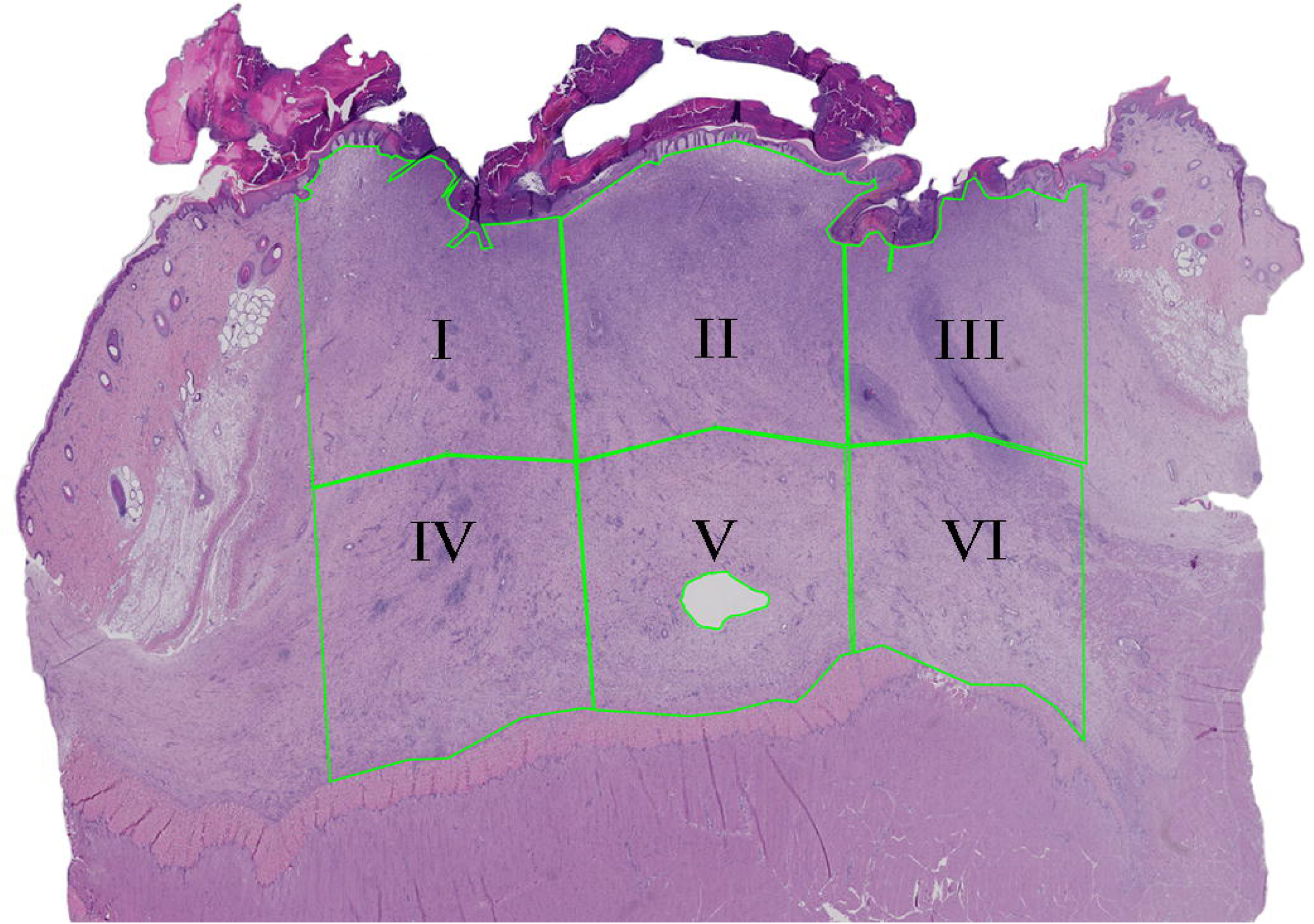
Hematoxylin and eosin-stained tissue from a 10-day old wound. The granulation tissue was divided into six approximately equal areas (I-VI) before histological evaluation

The epithelialized and non-epithelialized surface of granulation tissue was measured using the polyline tool in QuPath [30]. In each wound, the presence or absence of edema, accumulation of neutrophils in the most superficial part of the granulation tissue, and hemosiderophages were recorded. In each wound area, accumulation of leukocytes in the form of focal suppuration, pyogranulomas, or granulomas was recorded as present or absent.

### Assessment of granulation tissue maturation

A total of 50 wounds were randomly selected using the RANDBETWEEN function in Excel (Microsoft 365, Microsoft Corporation, Washington, USA). From each of these selected tissue sections, three images were taken to represent areas of granulation tissue (1000×525 μm) in the superficial, middle, and deep parts. The images were captured along the midline of the wound. All images (n=150) were grouped by Observer 1 as a) immature, b) established, or c) mature. Immature granulation tissue was characterized by a very high cellular density consisting of largecells, and blood vessels with either no visible lumen or a small diameter. Established granulation tissue was identified by the presence of parallel blood vessels, oriented perpendicular to the wound surface, with fibroblasts also aligned perpendicularly to the vessels. Mature granulation tissue was defined by blood vessels running parallel to the wound surface, a low cellular density, vessels with large diameters, and spindle-shaped cells. The images were then evaluated by Observer 2, who was blinded to both wound age and the depth of the tissue. Approximately two months later, the order of the images was randomized and re-evaluated by Observer 1, who was blinded to both wound age and the depth of the tissue. Then granulation tissue maturation was evaluated in each wound area in all wounds. As granulation tissue in a single wound area could consist of, e.g. both immature and established granulation tissue, the following classifications were used: 1) immature, 2) immature to established, 3) established, 4) established to mature, and 5) mature.

### Flow cytometry

All samples (control skin and granulation tissue) were washed in 1 mL phosphate buffered saline (PBS, pH 7.1-7.5) (Sigma-Aldrich, Catalog No. D8537) three times. The tissue was minced with a scalpel and was added to an Eppendorf tube containing PBS and 145 U/ml collagenase type I (Gibco, Thermo Fisher Scientific, Catalog No. 17100017) before incubation at 37°C for 2 hours while spinning in a MacsMix Tube Rotator (Miltenyi Biotec). The cell suspension was obtained by passing the digested tissue through a 70 µm strainer (Sarstedt AG & Co. KG, Order No. 83.3945.070) using 10 mL cold PBS. The cells were washed using cold PBS before measuring cell concentration to ensure sufficient cell count for analysis. The cells were washed and resuspended in 1 mL cold PBS. Then 1 µl live-dead stain (Table 1) was added and incubated for 30 minutes in dark, on ice. The cells were centrifuged, resuspended in cold PBS, and transferred to a 96-well plate. All cells were washed using cold PBS with 2% porcine serum (Gibco, Thermo Fisher Scientific, Catalog No. 26250084) prior to staining for CD34, CD45 and CD105 and their isotype controls (APC-Cy7-IgG, IgG1 and PE-IgG2a (Table 1)). Additionally, fluorescence minus one (FMO) controls were made for all antibodies. Antibody stains were incubated for 15 minutes in dark, on ice. Then 150 µl cold PBS was added to each well before centrifugation. Cells were resuspended in 150 µl cold PBS before being ready for analysis on the MACSQuant Analyzer 16 (Miltenyi Biotec).

**Table 1.**
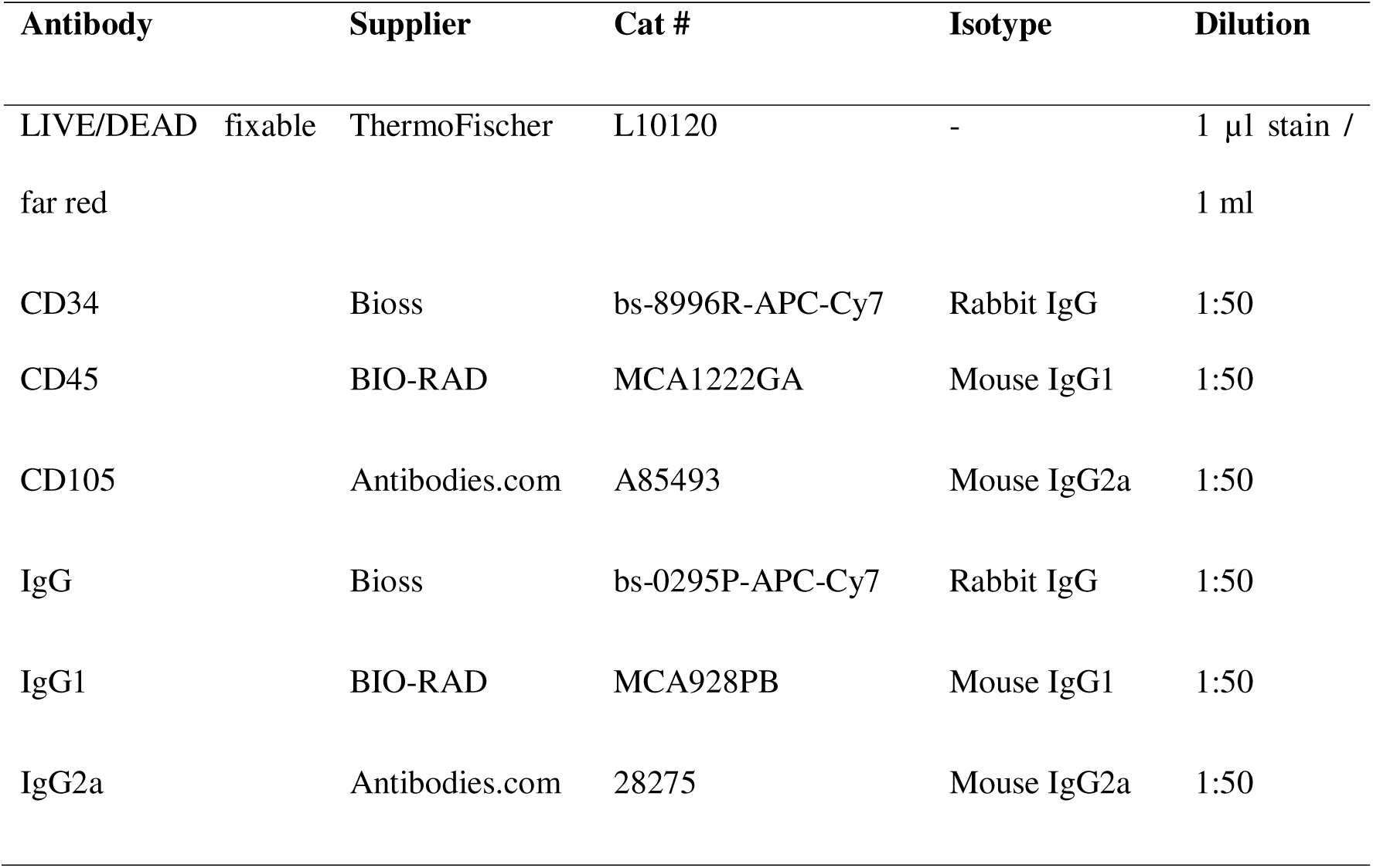
Antibodies used for flow cytometry, place of purchase and catalogue number, isotype of the antibody and the dilution used in the experiment.

### Immunohistochemical staining for CD45

Immunohistochemical staining was performed using the mouse anti-pig CD45 antibody (clone MAC323, IgG2a, Bio-Rad, MCA1447). Tissue sections (4-5 μm) were cut from the paraffin-embedded tissue blocks containing wound A from all pigs and intact skin from five pigs and a porcine lymph node (positive control). The sections were mounted on glass slides, deparaffinized, and rinsed in distilled water. Antigen retrieval was performed using TEG buffer at pH 8, followed by a cooling period in the same buffer. Sections were washed twice in Tris-buffered saline (TBS, pH 7.6) for 5 minutes. Endogenous peroxidase activity was blocked using 0.6% hydrogen peroxide in TBS for 15 minutes, followed by two additional washes in TBS for 5 minutes each. UltraVision Protein Block (TA-125-PBQ; Epredia, https://epredia.com) was applied for 5 minutes. The primary antibody was diluted 1:500 in 1% BSA/TBS and incubated overnight at 4°C. A nonspecific control (IgG2a, X0943; Agilent) was also applied at 1:500 in 1% BSA/TBS and incubated under the same conditions. After incubation, sections were washed in TBS twice for 5 minutes. Primary Antibody Enhancer (TL-125-PB) was applied for 20 minutes, followed by two washes in TBS for 5 minutes each. The UltraVision large volume HRP polymer (TL-125PH) was then applied for 30 minutes, followed by another two washes in TBS for 5 minutes each. Color development was performed using AEC (SK-4200; Vector Laboratories; https://vectorlabs.com) for 10 minutes. Sections were rinsed in running tap water twice for 5 minutes. Counterstaining was performed with Mayer’s hematoxylin for 30 seconds, followed by rinsing in distilled water twice for 5 minutes. Finally, sections were mounted using glycerol-gelatine.

### Positive cell detection

Tissue sections were digitized using an Axioscan 7 with a 20x/0.8 objective (Zeiss, Germany). Scanned tissue sections were imported into QuPath v0.5.1 [30], and the image type was set to Brightfield H-DAB and color deconvolution was performed. CD45 positive and negative cells were detected in all wound areas (I-VI) and control skin by running the Positive cell detection command (Table 2) [30]. To validate the detection of positive and negative cells, one wound was selected from each age group, along with one section of control skin. Within each selected wound, a wound area was identified, and a 300 × 300 µm annotation was placed at the center of this area. Wounds and wound areas were selected randomly using the RANDBETWEEN function in Excel. In the skin sample, the annotation was placed at the junction between the dermis and subcutis. True positives, false positives, true negatives, and false negatives were manually counted.

**Table 2.**
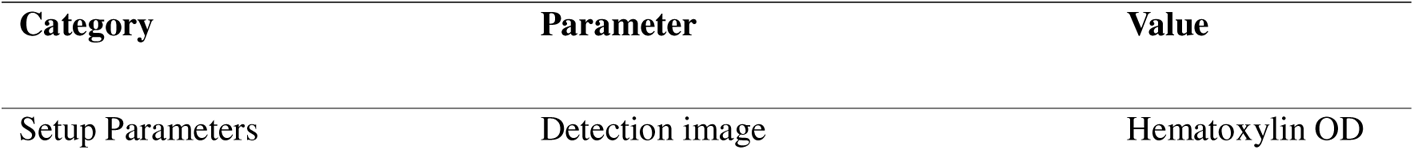
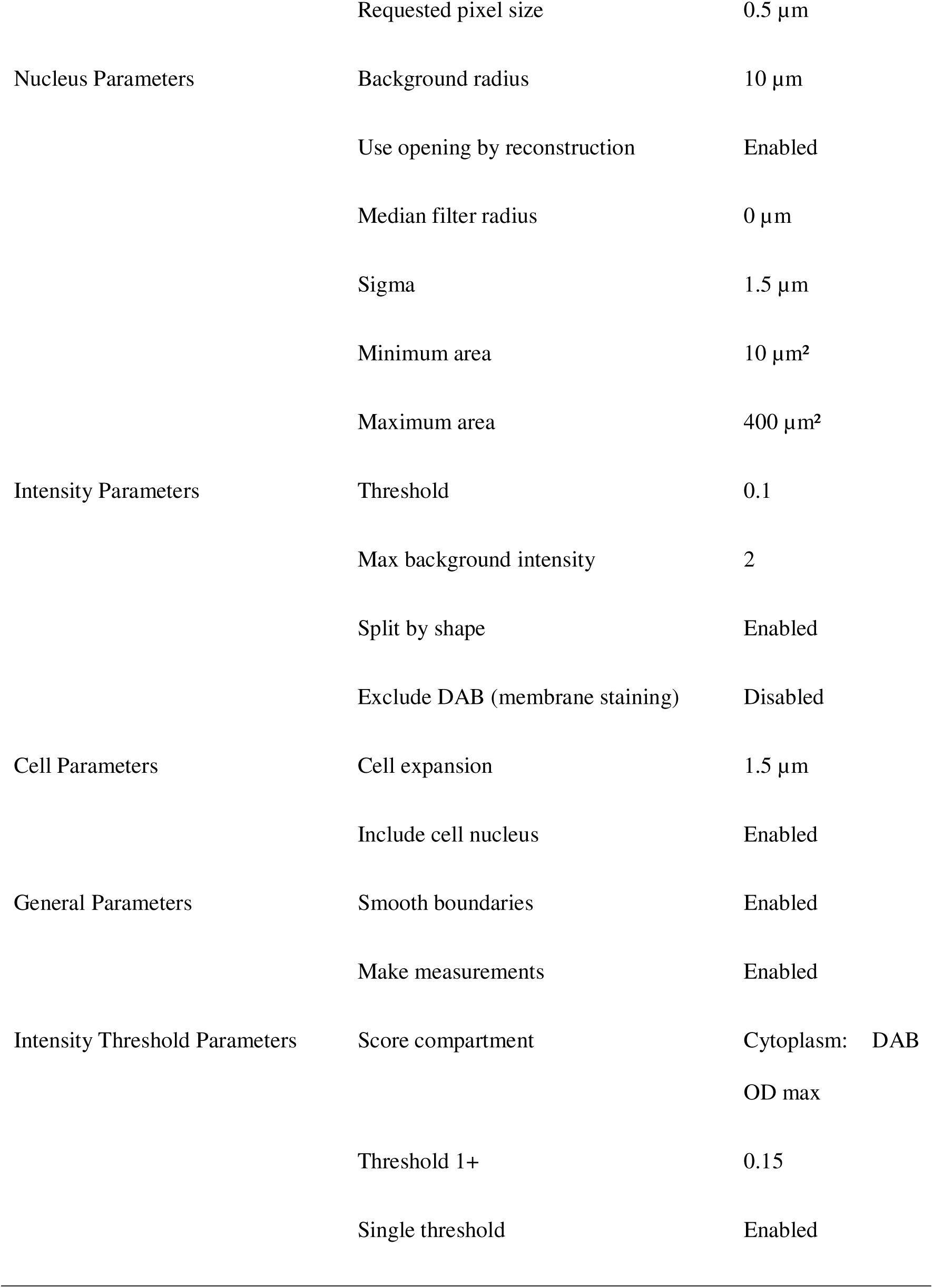
An overview of the different settings, and their parameters and values used for the positive cell detection in QuPath [30].

### Statistical analysis

Statistical analyses were conducted using Microsoft Excel, GraphPad Prism (version 10.4.2, GraphPad Software, San Diego, CA, USA) and RStudio (version 2024.12.0). Descriptive statistics were initially performed in Excel to summarize the dataset.

Weighted kappa was calculated using RStudio to assess inter- and intra-observer agreement for granulation tissue maturation. The strength of agreement was interpreted based on Landis and Koch criteria [31].

For flow cytometry analysis, dead cells were excluded using the live/dead stain. Living cells were used for further analysis. Isotype controls were used to set the positive stain limit at 1%. This gate was applied to the fully stained sample. FMO controls were used to verify that no background staining was present. One-sample t-test was performed in RStudio to test for statistically significant differences between wounds A and D.

For group comparisons, a one-way analysis of variance was conducted in GraphPad Prism to evaluate differences in the percentage of cells expressing CD34, CD45, and CD105 among the 8 groups (seven age groups and one control group). Tukey’s multiple comparisons test was applied to identify specific group differences. A significance level of α = 0.05 was used for all statistical tests.

## Results

### Gross evaluation

Granulation tissue was present in all wounds (n=188), and the tissue was thicker at the edges, compared to the center, in all groups (Fig. 3).

**Fig. 3.**
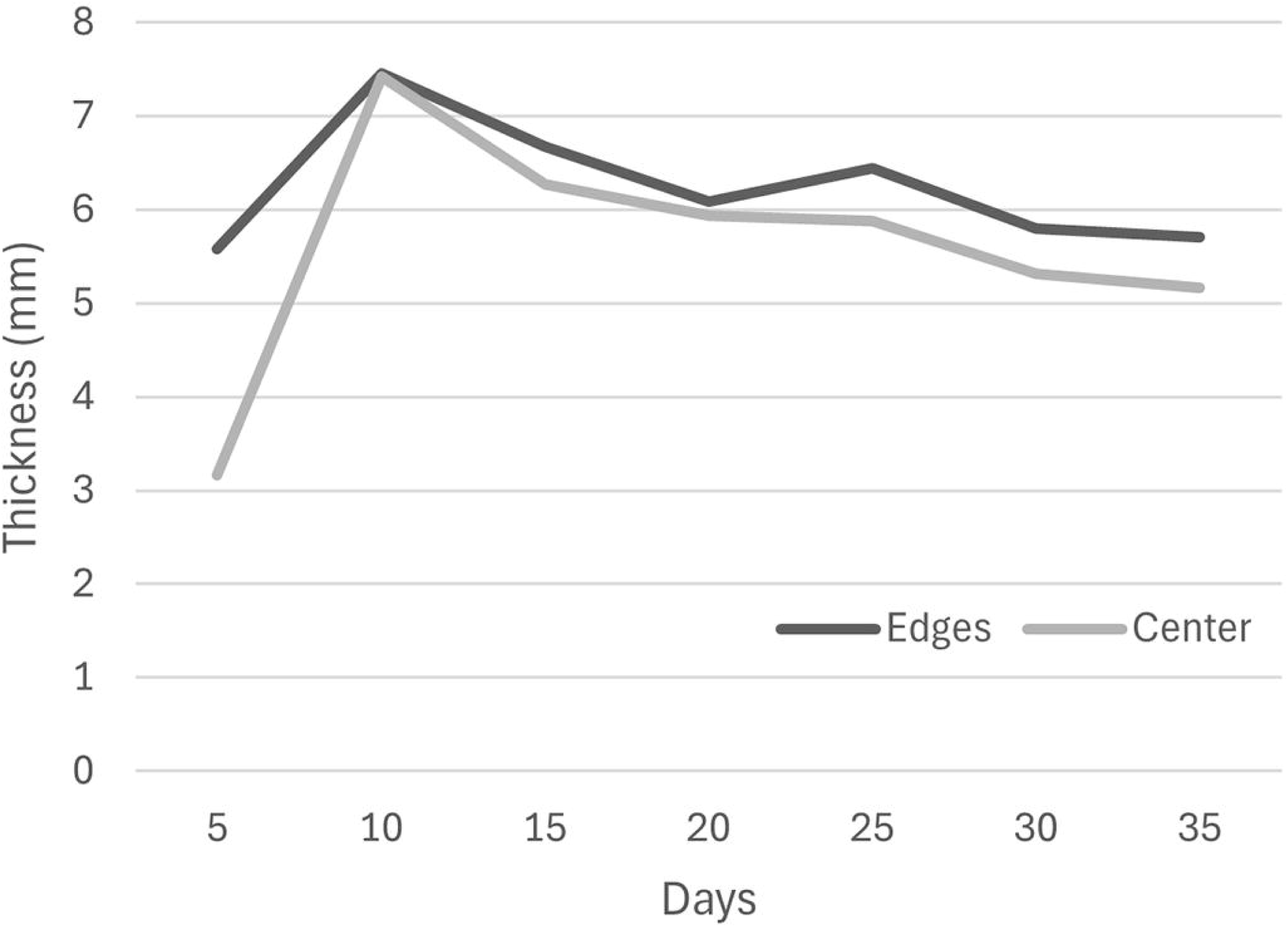
The thickness of the granulation tissue was measured postmortem. The thickness was measured at both the center and edges of the wounds. The values for each time point represent the average for all wounds of the given age

### Histological evaluation

A total of 179 wounds (1074 wound areas) were included, and nine wounds were discarded due to artefacts or sampling error (Table 3). In the 179 wounds included, a total of 1059 wound areas were suitable for histological evaluation, and 15 wound areas were discarded due to artefacts (Table 3). In addition, control skin, including epidermis, dermis, and subcutis, was included from 13 pigs.

**Table 3.**
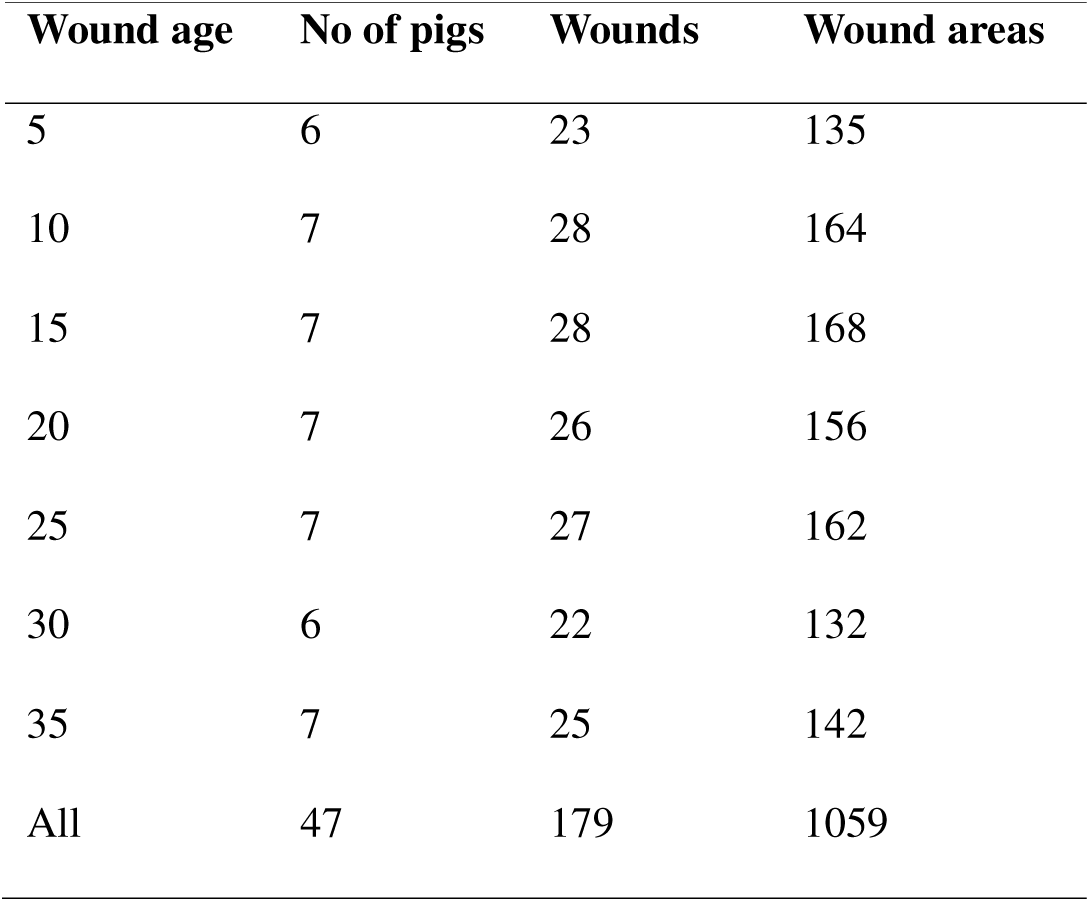
Wound age at time of euthanasia, number of pigs in each group, and number of wounds and wound areas included in the histological evaluations.

| Wound age | No of pigs | Wounds | Wound areas |
| --- | --- | --- | --- |
| 5 | 6 | 23 | 135 |
| 10 | 7 | 28 | 164 |
| 15 | 7 | 28 | 168 |
| 20 | 7 | 26 | 156 |
| 25 | 7 | 27 | 162 |
| 30 | 6 | 22 | 132 |
| 35 | 7 | 25 | 142 |
| All | 47 | 179 | 1059 |

Complete epithelialization was observed at the earliest in 20-day old wounds. The percentage of wound surface covered by epithelium increased with wound age (Fig. 4A). Edema was present in all 5-day old wounds, and the prevalence decreased as wound age increased. Edema was not present on day 35.

**Fig. 4.**
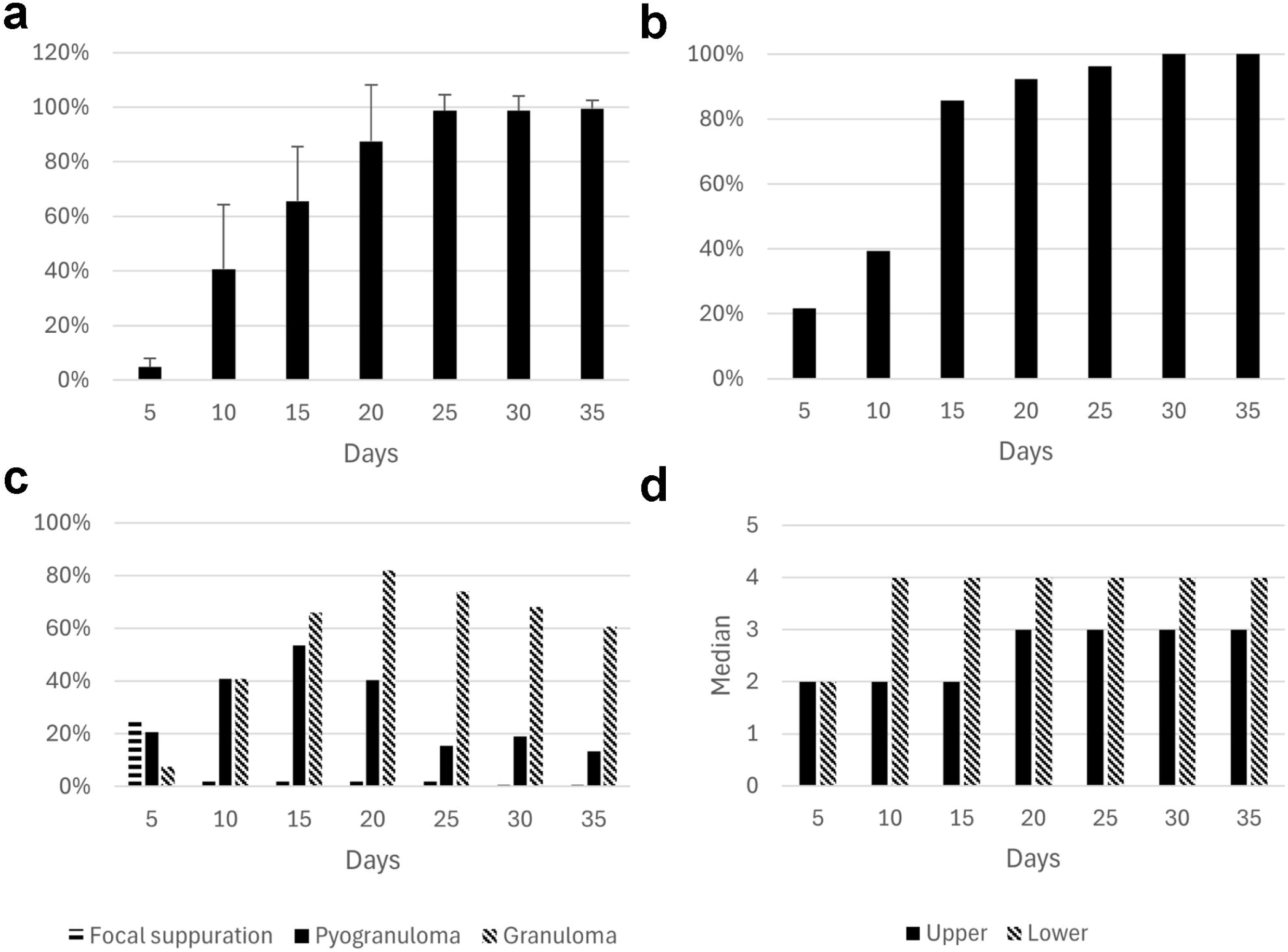
Histological evaluation of experimental porcine skin wounds. **a)** The epithelialized surface of granulation tissue expressed as the percentage (mean + standard deviation, SD) of the total surface of granulation tissue for each time point (n=179 wounds). **b)** The percentage of wounds (n=179 wounds) with presence of infiltrating hemosiderophages. **c)** Percentage of wound areas presenting with accumulation of leukocytes in the form of focal suppuration, pyogranulomas, or granulomas (n=1059 wound areas). **d)** Assessment of the maturation of granulation tissue in the superficial wound areas (I-III) and deep wound areas (IV-VI), respectively. In each wound area (n=1029) granulation tissue was classified as 1: immature, 2: immature to established, 3: established, 4: established to mature, or 5: mature. The median for each timepoint is presented in the histogram

Accumulations of neutrophils in the most superficial part of the granulation tissue (demarcation) were present in 100.0%, 92.8%, and 96.4% of wounds of 5, 10, and 15 days of age, respectively. Moreover, demarcation was seen in 34.6%, 7.4%, 0.0%, and 4.0% of wounds 20, 25, 30, and 35 days of age, respectively.

Hemosiderophages were observed at the earliest in wounds of 5 days of age and present in all 30- and 35-day old wounds (Fig. 4B).

Accumulations of leukocytes in the granulation tissue were present in all age groups (Fig. 4C). Focal suppuration was dominating in 5-day old wounds, while pyogranulomas peaked on day 15, and granulomas were predominantly seen from day 20 (Fig. 4C). Foreign bodies such as hair and plant material were seen regularly within granulomas and pyogranulomas.

No lesions were present in the control skin.

### Assessment of granulation tissue maturation

Substantial interobserver agreement (κ=0.719) and almost perfect intraobserver agreement (κ=0.920) were found for classification of granulation tissue as either immature, established, or mature (Tables 4 and 5). Confidence intervals were 0.606-0.832 and 0.876-0.964, respectively.

**Table 4.** Interobserver agreement based on granulation tissue maturation assessments performed by two veterinary pathologists.

|  |  | Observer 2 |  |  |
| --- | --- | --- | --- | --- |
|  |  | Immature | Established | Mature |
| Observer 1 | Immature | 23 | 4 | 0 |
|  | Established | 6 | 48 | 10 |
|  | Mature | 5 | 7 | 47 |

**Table 5.** Intraobserver agreement based on granulation tissue maturation assessment performed twice by the same veterinary pathologist, two months apart.

|  |  | Observer 1<sup>2nd time</sup> |  |  |
| --- | --- | --- | --- | --- |
|  |  | Immature | Established | Mature |
|  | Immature | 27 | 0 | 0 |
| Observer 1 | **Established** | 4 | 57 | 3 |
|  | **Mature** | 0 | 6 | 53 |

Granulation tissue was classified as immature (n=31 wound areas), immature to established (n =240 wound areas), established (n=322 wound areas), established to mature (n=387 wound areas), mature (n=49 wound areas), and unclassified (n=30 wound areas). The maturity of the granulation tissue increased with increasing wound age (Fig. 4D).

### Immunohistochemical IHC evaluation

A total of 39 wounds were included. Eight wounds were discarded due to artefacts or unsuccessful IHC staining. The 39 wounds consisted of 234 wound areas (I-VI), of which four were discarded due to artefacts. Control skin was included from five pigs.

Neutrophils, mononuclear cells, giant cells, and some spindle-shaped cells stained positive for CD45. Endothelial cells and most spindle-shaped cells did not express CD45 (Figs. 5A&B)

**Fig. 5.**
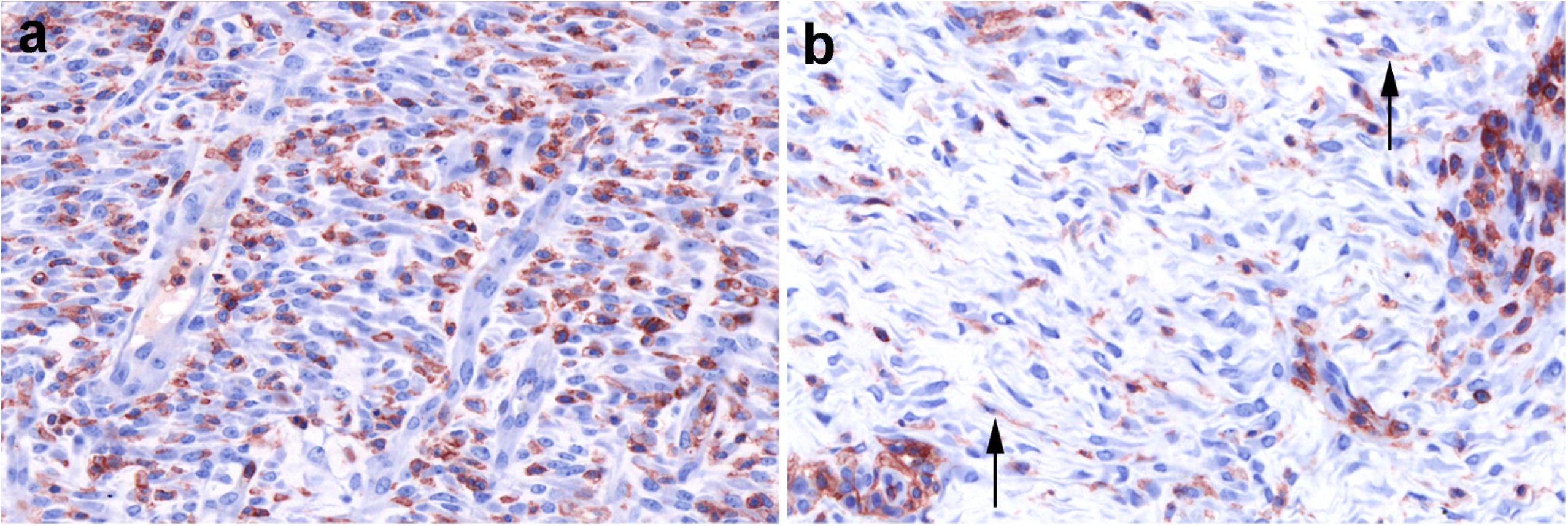
Immunohistochemical evaluation of experimental porcine skin wounds. **a)** Granulation tissue from a 5-day old wound immunohistochemically stained for cluster of differentiation 45 (CD45). Leukocytes and few spindle-shaped cells stain positive for CD45. **b)** Granulation tissue from a 30-day old wound immunohistochemically stained for CD45. Leukocytes and few spindle-shaped cells (arrows) stain positive for CD45

The positive cell detection algorithm was found to have a sensitivity of 89.7% (CI: 88.1-91.1%) and a specificity of 93.2% (CI: 92.0-94.3%). The validation was based on a total of 3298 cells categorized as true positives (n=1357 cells), false positives (n=121 cells), true negatives (n=1664 cells), and false negatives (n=156 cells).

In the superficial layer of the granulation tissue (wound areas I-III) the percentage of CD45 positive cells decreased as wound age increased, and a statistically significant difference was found between the groups (P<0.0001) (Fig. 6A). In the deep layer of the granulation tissue (wound areas IV-VI), the percentage of positive cells did not differ significantly between age groups (Fig. 6B).

**Fig. 6.**
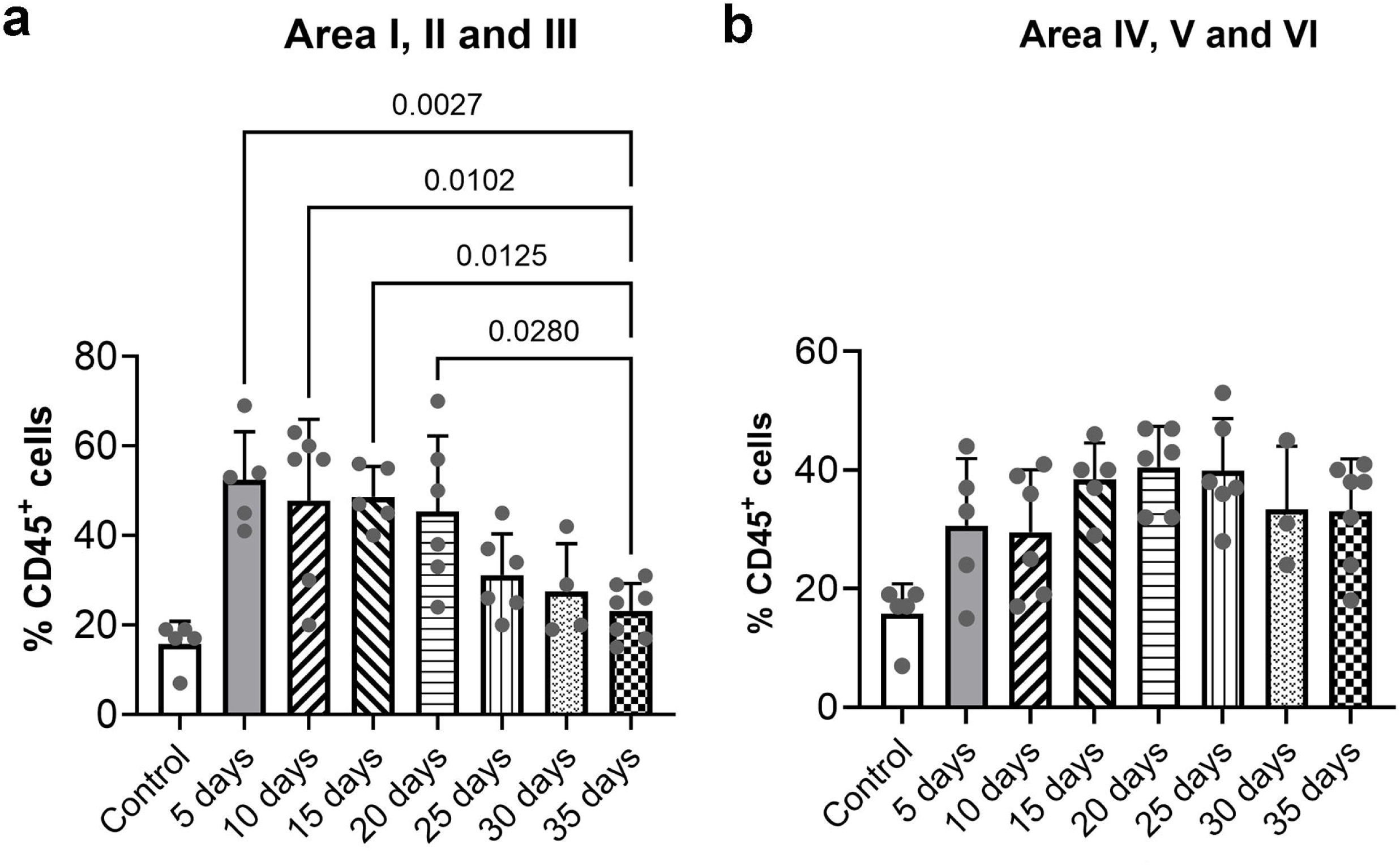
Percentages (mean + standard deviation, SD) of cells immunohistochemically positive for cluster of differentiation 45 (CD45). **a)** CD45 positive cells in superficial part of the granulation tissue (wound areas I-III) and **b)** CD45 positive cells in the deep part of the granulation tissue (wound areas IV-VI). Wound age ranges from 5-35 days and control samples are intact skin. A total of 39 wounds were included. One-way ANOVA with multiple comparisons was performed (P < 0.05)

### Flow cytometry

Wounds from 14 pigs were used for the development of the protocol and had to be excluded from the final analysis. Wounds from 33 of the pigs were included in the analysis for CD34. Furthermore, wounds from 36 pigs were included for CD105.

An average of wounds A and D was used since they were not statistically significantly different. Percentages of CD34 positive cells ranged from 7.45% to 25.02% (Fig. 7A), but there were no statistically significant differences between the groups. Percentages of cells positive for CD105 ranged from 0.80% to 11.63% (Fig. 7B) and the highest expression was observed in the 15-day old wounds. There was a statistically significant difference between wounds of 15 days of age and those of 30 and 35 days of age (p-values=0.0016 and 0.0027, respectively).

**Fig. 7.**
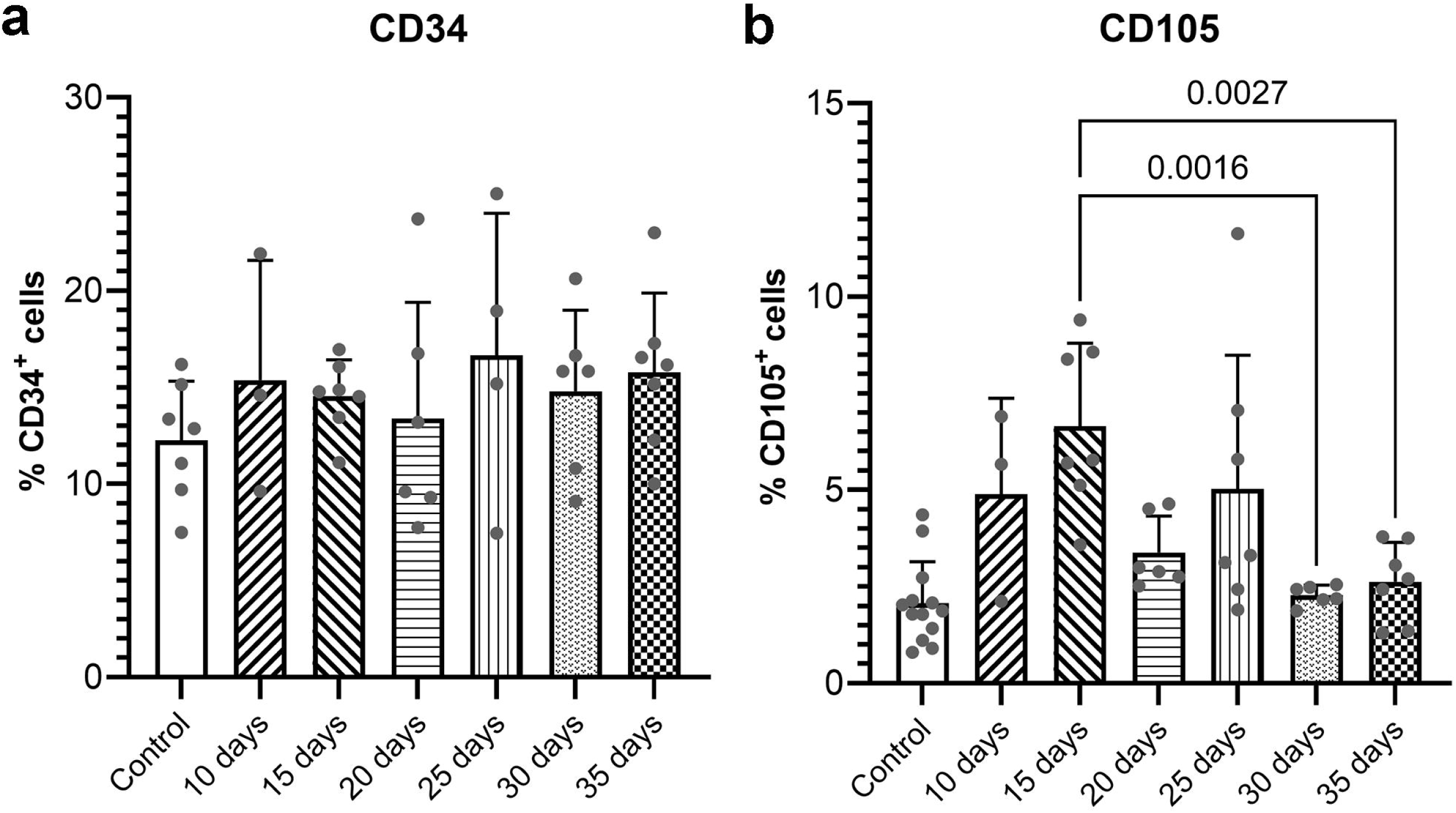
Flow cytometry histograms showing the percentage (mean + standard deviation, SD) of cells expressing: **A)** cluster of differentiation 34 (CD34), and **B)** cluster of differentiation 105 (CD105). Wound age ranges from 10-35 days and control samples are intact skin. A total of 33 and 36 wounds were included for CD34 and CD105, respectively. One-way ANOVA with multiple comparisons was performed and statistically significant differences are shown with P-values (P < 0.05)

## Discussion

In the present study the traditional methods of wound age assessments, macroscopic and histological evaluations, were applied to experimental porcine wounds in combination with IHC, digital pathology, and flow cytometry. Granulation tissue was macroscopically present in wounds from day 5 to 35. This is in accordance with previous studies in which granulation tissue was macroscopically visible in wounds from 5 days of age [5, 7, 32]. The thickness of the granulation tissue peaked at day 10, with a subsequent gradual decrease throughout the wound healing process (Fig. 3). This pattern was also reported in a previous porcine, experimental wound study [7] and underlines the importance that granulation tissue thickness alone is not suitable for age assessment. Histological assessment is to some extent subjective because the interpretation of the lesions may depend on the experience of the observer. However, in the present study interobserver agreement was found to be substantial (κ=0.719) for two veterinary pathologists assessing the maturity of granulation tissue. Furthermore, the granulation tissue classifications performed by a single veterinary pathologist, were found to be of almost perfect intraobserver agreement (κ=0.920). This highlights the reliability and usefulness of histological evaluations when performed by an experienced veterinary pathologist. However, the development of more objective methods remains necessary. Based on assessment of granulation tissue maturation, it was obvious that the lower part of the granulation tissue was classified as older, compared to the upper part of the granulation tissue. Knowing that wounds healing by second intention heal from the wound bed and upwards these findings were expected [33].

Re-epithelialization is considered to be among the most important steps of wound healing [34, 35]. We observed that epithelialization had started in the 5-day old wounds. In comparison, previous studies in humans, pigs, and mice have shown that epithelialization begins 16-24 hours post wounding [36–38]. When the skin barrier is broken upon injury, leukocytes are recruited to the site of injury to help kill microbes, clear debris, and protect the wound environment until the epithelium is reestablished [39, 40]. Accumulation of leukocytes was seen in all age groups. Focal suppuration was primarily seen in the 5-day old wounds. Granulomas peaked in the 20-day old wounds, while pyogranulomas were the highest in 15-day old wounds. This is in accordance with the study by Barington et al, 2018 [7]. In that study, neutrophils dominated in the early experimental wounds while macrophages were the dominant cell type in the wounds from day 14.

Hemosiderophages were found to be present at all wound ages. However, they were only present in all wounds of 30 and 35 days of age. Barington et al, 2018 [7] similarly found hemosiderophages present from 8 to 35 days post wounding in an excisional wound model. These findings correspond with what has been observed in humans [41].

Fibrocytes, which are CD34 positive, have been shown to lose the CD34 marker upon differentiation to myofibroblasts [42, 43]. We targeted CD34 for its potential to be expressed in a time-dependent manner since myofibroblasts play a role in wound contraction [44]. However, in the present study no time-dependent pattern in the percentage of CD34 positive cells was seen. A time-dependent expression of CD105 was observed with a peak on day 15. CD105 is a known marker of endothelial cells involved in the formation of new blood vessels [21–23]. In comparison, Pankoke et al [5] showed that vessel density in granulation tissue peaked on day 6 post wounding, with a subsequent gradual decrease during healing. We did see lower percentages of CD105 positive cells compared to previous studies using this marker to assess cells in granulation tissue from intra-bony periodontal defects in humans [45, 46]. However, it is possible that the cells positive for these markers are not as readily available for extraction from porcine granulation tissue [47] as these tissues consist of dense connective tissue and extracellular matrix [48].

CD45 is a well-known marker of HCs, which include both leukocytes and fibrocytes [20, 42, 49]. The high percentage of CD45 positive cells (Fig. 6) observed in all age groups of the experimental wounds, is suspected as leukocytes play a role during most wound healing phases. Moreover, few spindle shaped cells expressed CD45. These cells could potentially be fibrocytes, which are positive for CD45 [49]. A significant difference in CD45 positive cells was observed in the superficial part of the granulation tissue, however, there was no significant difference between the age groups in the deep part of the granulation tissue. Expression was found to be highest in the youngest wounds, with a decrease in percentage of positive cells, as wound age increased. Additionally, the sensitivity between the manual count performed by the veterinary pathologist and the digital counter was very high, showing that a cell detection algorithm can provide trustworthy data. The algorithm allows for objective, quantitative assessment of much larger areas than what is feasible to evaluate manually.

## Conclusion

During granulation tissue formation and maturation, time-dependent changes in several factors, including re-epithelialization, hemosiderophages, granulation tissue maturity classification, and expression of CD105 and CD45 were observed. Current age estimations based on granulation tissue thickness and histological evaluations are not sufficient for an accurate wound age assessment. The addition of flow cytometry and IHC combined with digital pathology may help to obtain a more objective and accurate age estimation.

## Key points

1. Substantial interobserver agreement was reached on the histological evaluations between two experienced veterinary pathologists.
2. Macroscopic and histological evaluations alone are not objective or precise enough for accurate age assessments.
3. A multidisciplinary approach with flow cytometry and IHC combined with digital pathology can help obtain more accurate and objective age estimates of wounds in veterinary forensic cases.

## Statement of author contributions

Conceptualization: Cecilie Bækgård, Henrik Elvang Jensen, Kristiane Barington; Methodology: Cecilie Bækgård, Lars Andresen, Marie Høy Hansen, Henrik Elvang Jensen, Kristine Freude, Kristiane Barington; Formal analysis and investigation: Cecilie Bækgård, Marie Høy Hansen, Henrik Elvang Jensen, Kristiane Barington; Writing – original draft preparation: Cecilie Bækgård; Writing – review and editing: Lars Andresen, Marie Høy Hansen, Henrik Elvang Jensen, Kristine Freude, Kristiane Barington; Funding acquisition: Kristiane Barington; Supervision: Kristiane Barington.

All authors read and approved the final manuscript. The manuscript has not been accepted or published elsewhere.

## Acknowledgements

The authors would like to thank Betina Gjedsted Andersen and Elisabeth Wairimu Petersen for assistance with preparation of tissue for histology and immunohistochemistry. We also acknowledge Dennis Brok and Frederik Andersen for technical assistance.

## Statements and Declarations Ethics approval

The study and procedures were approved by the Danish Animal Inspectorate (2023-15-0201-01363).

